# A transient epithelial plasticity state defines the developmental window for uterine gland specification

**DOI:** 10.64898/2026.06.03.729801

**Authors:** Jason A. Rizo, Abdallah W. Abdelhady, Valentina Lorenzi, Dhanvika Mopure, Jacob M. Pru, Sarayut Winuthayanon, Roser Vento-Tormo, Ciro M. Amato, Thomas E. Spencer, Andrew M. Kelleher

## Abstract

Uterine gland development and function is essential for reproduction and women’s health, yet the epithelial cell states and signaling interactions that govern gland fate specification are not well understood. Here, integration of single cell and spatial transcriptomics with organoid culture, lineage tracing, and genetic and hormonal perturbation models were used to define mechanisms regulating postnatal uterine epithelial differentiation. A developmentally restricted epithelial plasticity state was identified that precedes luminal and glandular cell lineage segregation and is accompanied by dynamic reorganization of stromal-epithelial communication during uterine differentiation. Pseudotime analysis revealed progressive acquisition of gland-associated programs, including *forkhead box A2* (*Foxa2*), retinoic acid metabolic genes, and epithelial *estrogen receptor alpha* (*Esr1*) expression. Functional studies revealed that ESR1 acquisition and retinoic acid signaling suppress the multilayered organoid phenotype associated with epithelial plasticity, thereby promoting epithelial specification and lineage commitment. Moreover, neonatal hormonal perturbation of adenogenesis and conditional deletion of *Foxa2* abolished this organoid phenotype. Together, these findings demonstrate that ESR1 acquisition, retinoic acid signaling and FOXA2-dependent glandular differentiation each restrict a transient epithelial plasticity state, coupling the loss of developmental plasticity to the emergence of the glandular lineage.

## INTRODUCTION

Uterine development and function depend on tightly regulated paracrine interactions and hormonal signaling that establish and maintain tissue homeostasis^1,2^. During development, the paired Müllerian ducts undergo coordinated morphogenesis and differentiation to form the structure of the adult uterus, including the epithelium, stroma, endothelium, and myometrium^3,4^. Genetic and environmental perturbations during critical developmental windows can disrupt these processes, leading to congenital anomalies, impaired reproductive function, and increased susceptibility to gynecologic disease, including endometriosis, uterine squamous metaplasia, and uterine cancer^5–10^. Despite the importance of these processes for reproductive health, the cellular and molecular pathways that govern uterine development remain incompletely understood.

Single-cell and spatial transcriptomic studies have begun to resolve the cellular heterogeneity and developmental trajectories of the developing female reproductive tract (FRT)^11–14^. These analyses indicate that uterine epithelial maturation is governed by both intrinsic gene regulatory programs as well as inductive and instructive cues from the surrounding mesenchyme and stroma. Although human and mouse uterine anatomy and the timing of gland morphogenesis or adenogenesis differ, major uterine cell types and many signalling pathways are broadly conserved, supporting the use of mouse models to define core mechanisms of uterine morphogenesis^11–14^. This comparative framework is especially important because access to human fetal and neonatal FRT tissues is inherently limited, precluding direct analysis of developmental states in the uterine epithelium, and *in vitro* systems that faithfully recapitulate Müllerian epithelial development from pluripotent stem cells remain underdeveloped. In both mice and humans, uterine morphogenesis involves stromal patterning, myometrial expansion, vascular organization, and the initiation of adenogenesis^3,4,15–17^.

Classic tissue recombination studies established that the FRT epithelium loses competence to respond to heterotypic mesenchyme during the first week of postnatal life^18–21^. More recently, organoid and assembloid models have enabled direct interrogation of this inherent epithelial plasticity and modelling of epithelial-mesenchymal crosstalk *in vitro*^22,23^. Those studies revealed that the neonatal uterine epithelium transiently retains developmental plasticity and remains responsive to instructive cues from the mesenchyme/stroma during the early postnatal period^22^. In mice, the gradual developmental restriction of epithelial plasticity in the uterus coincides with the timing of epithelial lineage specification and the initiation of adenogenes^18–21^. However, the cellular and molecular mechanisms that restrict cellular plasticity, the cell states that precede commitment of epithelial lineages, and the signaling interactions governing stroma-regulated luminal and glandular cell fate specification in the endometrium remain to be established^24–26^.

Adenogenesis is a critical event in FRT development as defects in uterine glands compromise fertility and are associated with uterine pathologies, emphasizing the clinical relevance of studying uterine development to understand disease^27–30^. Prior studies have implicated steroid hormone signaling, retinoic acid metabolism, and FOXA2-dependent gene regulatory networks in uterine epithelial differentiation and disease progression^31–34^. How these pathways impact transition from an immature epithelial cell state to committed luminal and glandular cell lineages remain unresolved. Whether these epithelial fate transitions can be captured during uterine development and disease progression, and functionally dissected in organoid culture, remains untested^22,35,36^.

Here, we generated a single-cell atlas of the fetal and neonatal mouse uterus to define the cellular states and signaling crosstalk that govern epithelial fate specification. By integrating multiomic approaches with genetic and hormonal mouse models of uterine gland ablation, lineage tracing, and organoid culture, these studies identify a developmentally transient epithelial state that precedes glandular cell differentiation, define dynamic stromal-epithelial signaling interactions during neonatal uterus maturation, and find that ESR1 acquisition and retinoic acid signaling restrict cellular plasticity and promote lineage commitment towards single-layered columnar epithelium. This transient cell state is captured *in vitro* by a multilayered organoid phenotype that is progressively lost during uterine development and abolished by hormonal and genetic disruption of adenogenesis, demonstrating that multiple developmental inputs act within a narrow postnatal window to couple the loss of epithelial plasticity to glandular differentiation.

## RESULTS

### Single-Cell Transcriptomic Atlas of the Developing Mouse Uterus

Single-cell RNA sequencing (scRNA-seq) was performed on uterine tissue collected at embryonic day 16.5 (E16.5) and postnatal days (PND) 1, 5, 12, and 15 to characterize the transcriptional dynamics of the developing mouse uterus at single cell resolution. Enzymatically dissociated uterine cells were subjected to EpCAM-based magnetic sorting to achieve an approximately equal ratio of epithelial (EpCAM-positive) and non-epithelial (EpCAM-negative) cells prior to library preparation using the 10X Chromium Flex system (Fig. 1A). After quality control and filtering (see Methods), a total of 147,440 cells were retained for downstream analysis. Unsupervised clustering resolved 16 transcriptionally distinct populations, annotated into 8 major cell lineages based on canonical marker gene expression (Fig. 1B-D; Dataset 1). The epithelial (47,928 cells; *Cdh1*, *Epcam*, *Tacstd2*) and stromal (72,561 cells; *Vim*, *Pdgfra*, *Hoxa10*) compartments comprised the two largest populations. The epithelial lineage resolved into three clusters corresponding to Müllerian duct epithelium (MDE) and two temporally staged populations (PND1-5; PND12-15), while the mesenchyme was resolved into one Müllerian duct ligament and two mesenchyme (MDM) clusters, two temporally staged populations (PND1-5; PND12-15), and uterine ligament (Fig. 1C-D; Sup. Fig. 1A-B). Additional lineages included myometrium (9,457 cells; *Myh11*, *Actg2, Acta2*), endothelium (7,295 cells; *Pecam1*, *Robo4*), coelomic epithelium (4,250 cells; *Muc16*, *Lrrn4*), perivascular (2,515 cells; *Rgs5*, *Notch3*), immune (2,837 cells; *Cd53, Tyrobp*), and neural (597 cells; *Tubb3, Phox2b*) (Fig. 1B-D).

**Figure 1.**
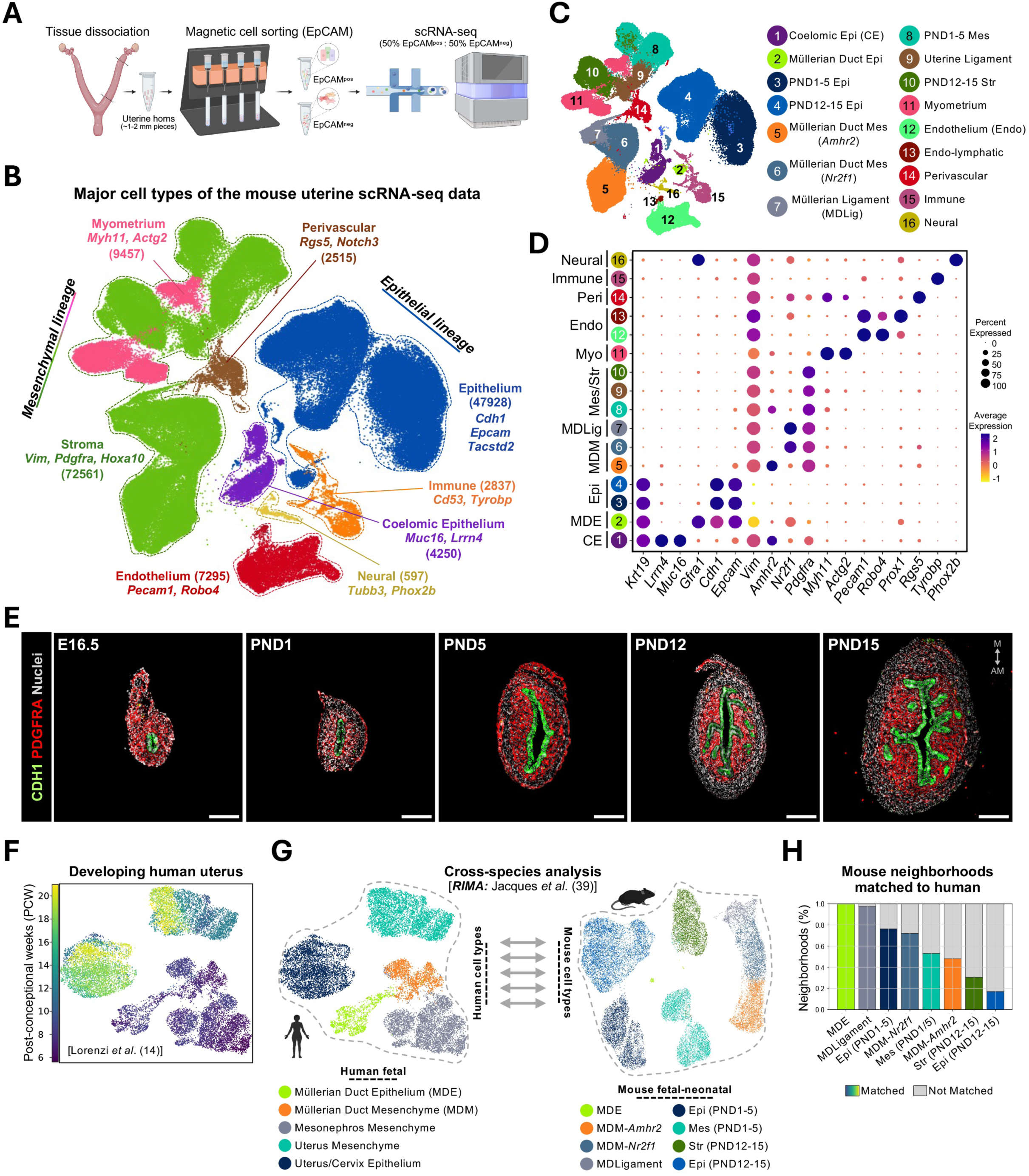
Single-cell transcriptomic atlas of the developing uterus. (A) Schematic of the experimental workflow. Uterine horns were collected at embryonic day 16.5 (E16.5) and postnatal days (PND) 1, 5, 12, and 15, enzymatically dissociated, and subjected to EpCAM-based magnetic cell sorting to achieve an approximately equal ratio of EpCAM-positive (epithelial) and EpCAM-negative (non-epithelial) cells prior to 10x Chromium Flex scRNA-seq library preparation. (B) Uniform manifold approximation and projection (UMAP) embedding of the scRNA-seq dataset (*n* = 147,440 cells) coloured by major developmental cell lineages. Cell numbers and defining marker genes are indicated for each population: epithelium (47,928; *Cdh1, Epcam, Tacstd2*), stroma (72,561; *Vim, Pdgfra, Hoxa10*), myometrium (9,457; *Myh11, Actg2, Acta2*), endothelium (7,295; *Pecam1, Robo4*), coelomic epithelium (4,250; *Muc16, Lrrn4*), immune (2,837; *Cd53, Tyrobp*), perivascular (2,515; *Rgs5, Notch3*), and neural cells (597; *Tubb3, Phox2b*). (C) UMAP from (B) coloured by cluster (left) and description of cell cluster identity (right). (D) Dot plot showing scaled average expression (color) and percentage of cells expressing (dot size) canonical marker genes across cell clusters. (E) Representative immunofluorescence images of CDH1 and PDGFRA in uterine cross-sections at E16.5, PND1, PND5, PND12, and PND15. Scale bars, (100 μM). (F) UMAP embedding of human fetal uterine epithelial and mesenchymal cells from Lorenzi *et al*.^14^. Uterine tissues were collected from post-conceptional week (PCW) 6 to 20 as indicated by color (n=18,243 cells). (G-H) Cross-species analysis of epithelial and mesenchymal lineages using RIMA^39^ (G) and visualization of the percentage of mouse neighborhood matched to human (H). Abbreviations: PND, post-natal day; MD, Müllerian duct; MDE, Müllerian duct epithelium; MDM, Müllerian duct mesenchyme; CE; coelomic epithelium; EPI, epithelium; MES, mesenchyme; STR, stroma; MYO, myometrium; Endothelium-L, endothelial-lymphatic; PERI; perivascular; M, mesometrial; AM, anti-mesometrial.

Visualization of cells by developmental timepoint revealed dynamic shifts in cluster composition, and substantial transcriptional changes occurring between E16.5 and PND15 (Figs. 1C-D; Sup. Figs. 1A–C). Notably, markers of smooth muscle (*Acta2*, *Actg2*, and *Myh11*) and endothelium (*Pecam1, Robo4*) were detected as early as E16.5 (Fig. 1C-D, and Sup. Fig. 1A-D). This early specification was validated at the protein level by immunofluorescence, revealing a thin but distinct ACTA2-positive muscle layer in the outer mesenchyme at E16.5 that expanded and became progressively organized between PND5 and PND15. Similarly, PECAM1-positive endothelial cells were present at E16.5 and formed nascent vascular networks, suggesting that smooth muscle and vascular lineage specification initiates prior to birth, with continued postnatal expansion and organization (Sup. Fig. 1E).

We then integrated our dataset with published scRNA-seq (GSE275806^37^) of the adult mouse uterus, using CellHint^38^ for label harmonization (Sup. Fig. 1F-G). The adult data included 19 populations representing 3 luminal epithelium (LE), 4 glandular epithelium (GE), 2 stromal, 3 endothelial, 2 perivascular, 2 mesothelial, and 3 immune cell clusters (Sup. Fig. 1F). CellHint analysis revealed that non-epithelial cell types including myometrial, endothelial, perivascular, and immune populations mapped to their adult counterparts, indicating broad conservation of these lineage identities across developmental stages. In contrast, epithelial and mesenchymal clusters from PND1 and PND5 were exclusive to the fetal-neonatal dataset, possibly representing developmentally restricted transcriptional cell states that do not persist into adulthood^37^. By PND12 and PND15, epithelial clusters began to align with adult luminal and glandular cell subtypes (LE_2, GE_1), coinciding with the onset of adenogenesis in mice (Fig. 1E). However, the more transcriptionally mature luminal (LE_1, LE_3) and glandular (GE_2, GE_3, GE_4) cell populations in the adult uterus remained unique, indicating that terminal epithelial differentiation occurs after the postnatal period between weaning and puberty (Sup. Fig. 1F-G).

This single-cell resource captures the transcriptional landscape of the developing mouse uterus across 8 major cell lineages and 5 developmental timepoints, providing a comparative framework for cross-species analysis of early uterine development. It covers the progression from undifferentiated Müllerian duct precursor cells through the initial emergence and establishment of luminal and glandular epithelial cell identities. The resource is accessible at www.genesearch.org/sklab/uterineatlas/.

### Cross-Species Comparison of Human and Mouse Uterine Development

To determine whether the transcriptional programs characterizing mouse fetal and neonatal uterine development are conserved in humans, the mouse data was subset to mesenchymal and epithelial cell populations (n=31,969) and compared to human Müllerian duct and uterine cells from post-conceptional weeks (PCW) 6 to 20^14^ (Fig. 1F-G). This analysis was performed using RIgorous Matching of Atlases (RIMA), a neighborhood-level matching approach that operates on species-specific low-dimensional embeddings without forcing cross-species integration, recently developed by Jacques *et al*.^39^ (Fig. 1G). Human fetal cell types (n=18,243) included MDE and MDM, mesonephros mesenchyme, uterine mesenchyme, and uterus/cervix epithelium (Fig. 1F-G; Sup. Fig. 2A-B). RIMA revealed that mouse cell neighborhoods from E16.5 through PND5 showed high concordance with their human counterparts, whereas PND12-15 neighborhoods were largely unmatched (Fig. 1H), which is likely due to the absence of adenogenesis in second-trimester human uterine samples reported by Lorenzi *et al*.^14^. We then leveraged the matched neighborhoods to identify genes with the most conserved and most divergent expression patterns across species within shared cell types (Sup. Fig. 2C-F). Within matched epithelial neighborhoods, the most conserved genes were enriched for transcriptional regulators of Müllerian duct identity including *PAX2, PAX8*, *SOX17*, *MSX1, MSX2*, and *WNT7A*, as well as core epithelial junction and polarity components including *CDH1*, *TACSTD2*, *KRT7*, *KRT18*, and *CLDN3*, suggesting that the transcriptional programs governing luminal epithelial identity in the pre-adenogenesis uterus are conserved between mouse and human fetal development^40–47^ (Fig. 1H; Sup. Fig. 2F). In contrast, the least conserved genes included *GAS6, SOCS2, SMOC2, ELOVL5, HSD11B2,* and *APOC1* representing stromal-derived signaling whose expression reorganizes alongside adenogenesis in mice, as well as species-specific metabolic and extracellular matrix programs, consistent with the absence of active adenogenesis in the human fetal samples during the first and second trimester of prenancy^14,48^ (Sup. Fig. 2E).

**Figure 2.**
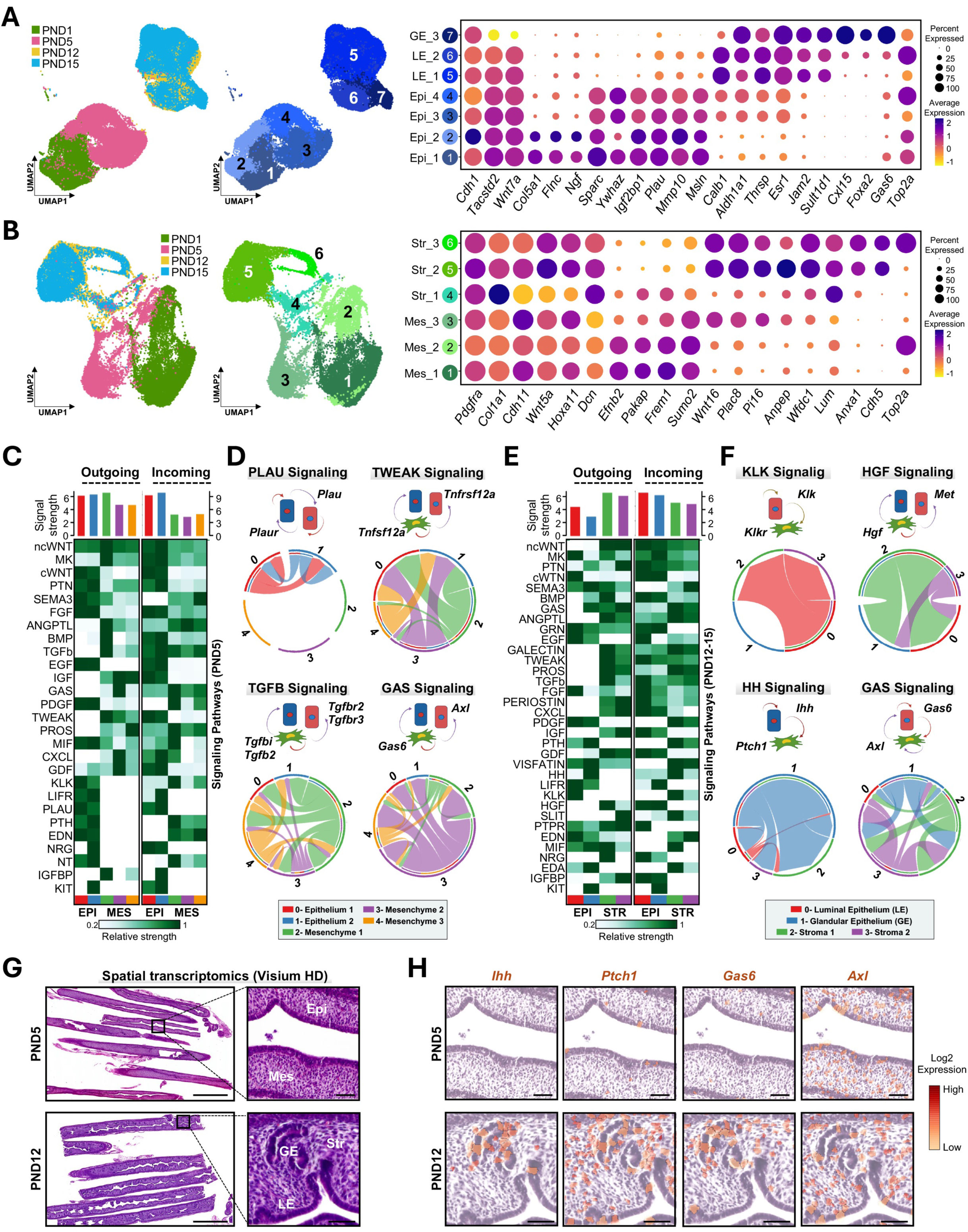
Epithelial and stromal cell identities and stromal-epithelial signaling dynamics during postnatal uterine development. (A) Uniform Manifold Approximation and Projection (UMAP) embedding of epithelial cells (n = 61,486) in postnatal scRNA-seq samples > PND1 coloured by developmental timepoint (left) or epithelial subcluster identity (middle). Dot plot (right) showing scaled average expression (color) and percentage of cells expressing (dot size) selected marker genes defining early epithelial clusters (Epi_1–4), luminal epithelial clusters (LE_1–2), and the glandular epithelium (GE_3). (B) UMAP embedding of mesenchymal/stromal cells (n = 30,251) coloured by developmental timepoint (left) or subcluster identity (middle). Dot plot (right) showing selected marker genes defining early mesenchymal clusters (Mes_1–3) and mature stromal clusters (Str_1–3). (C) Heatmap from CellChat analysis comparing the outgoing (left) and incoming (right) signaling patterns associated with both mesenchymal and epithelial cells at PND5. Shading denotes the relative signalling strength of a pathway across cell types. Coloured bar plots on top depict the signalling strength of a particular cell cluster by summarizing all pathways in the heatmap. (D) Schematic of ligand-receptor interaction networks at PND5 highlighting mesenchymal-epithelial crosstalk for selected signalling pathways. (E) Heatmap from CellChat analysis comparing the outgoing (left) and incoming (right) signaling patterns associated with both mesenchymal and epithelial cells at PND12-15. (F) Schematic of ligand-receptor interaction networks at PND12-15 highlighting stromal-epithelial crosstalk for selected signalling pathways. (G) Visium HD spatial transcriptomic analysis of longitudinal uterine sections from PND5 and PND12 (left, scale bar, 1 mm; right, insets 50 μm). (H) Spatial expression of selected genes at PND5 (top) and PND12 (bottom), showing reorganization of stromal-epithelial signalling during postnatal uterine maturation. Scale bars, 50 μm. Abbreviations: PND, post-natal day; EPI, epithelium; MES, mesenchyme; STR, stroma; LE, luminal epithelium; GE, glandular epithelium.

### Epithelial and Mesenchymal Cell Identities During Postnatal Uterine Development

Epithelial and stromal cells were subset and re-clustered across postnatal development (see Methods) to interrogate lineage specific subpopulations before and after the initiation of adenogenesis^11,16,17,27,49^ (Fig. 2A-B). Epithelial subclustering identified 7 transcriptionally distinct populations that reflected both developmental stage and lineage identity^11–13^ (Fig. 2A). Four early clusters (Epi_1-4) were enriched on PND1 and PND5 and expressed the luminal epithelial markers *Cdh1* and *Wnt7a*. Notably, these clusters expressed genes not associated with mature cell types including *Ywhaz*, *Plau,* and *Msln*, suggesting that the early postnatal epithelium retains a transcriptionally unspecified state prior to uterine fate commitment (Fig. 2A). In line with this, GSEA of PND5 epithelial clusters revealed enrichment for G2M Checkpoint and E2F target gene sets, indicating that the unspecified epithelium is characterized by a proliferative state (Sup. Fig. 3A). By PND12 and PND15, luminal (LE_1, LE_2) and glandular (GE_3) cell clusters emerged with shared markers including *Thrsp*, *Esr1*, *Jam2,* and *Sult1d1*, (Fig. 2A) and pathway enrichment for Estrogen Response hallmarks (Sup. Fig. 3A). The LE clusters were specified by *Calb1*, *Wnt7a*, and *Tacstd2*, while the GE was uniquely defined by *Foxa2*, *Gas6*, and *Cxcl15* (Fig. 2A). This lineage segregation was accompanied by specific enrichment for P53 Signaling, IL6-JAK-STAT3, and Apical Junction in the LE, while the GE cluster exhibited enrichment for NOTCH Signaling, Protein Secretion, and Oxidative Phosphorylation hallmarks, consistent with a specialized secretory identity (Sup. Fig. 3A). These findings suggest that the early postnatal epithelium represents a transcriptionally immature population that has not yet acquired luminal or glandular identity.

**Figure 3.**
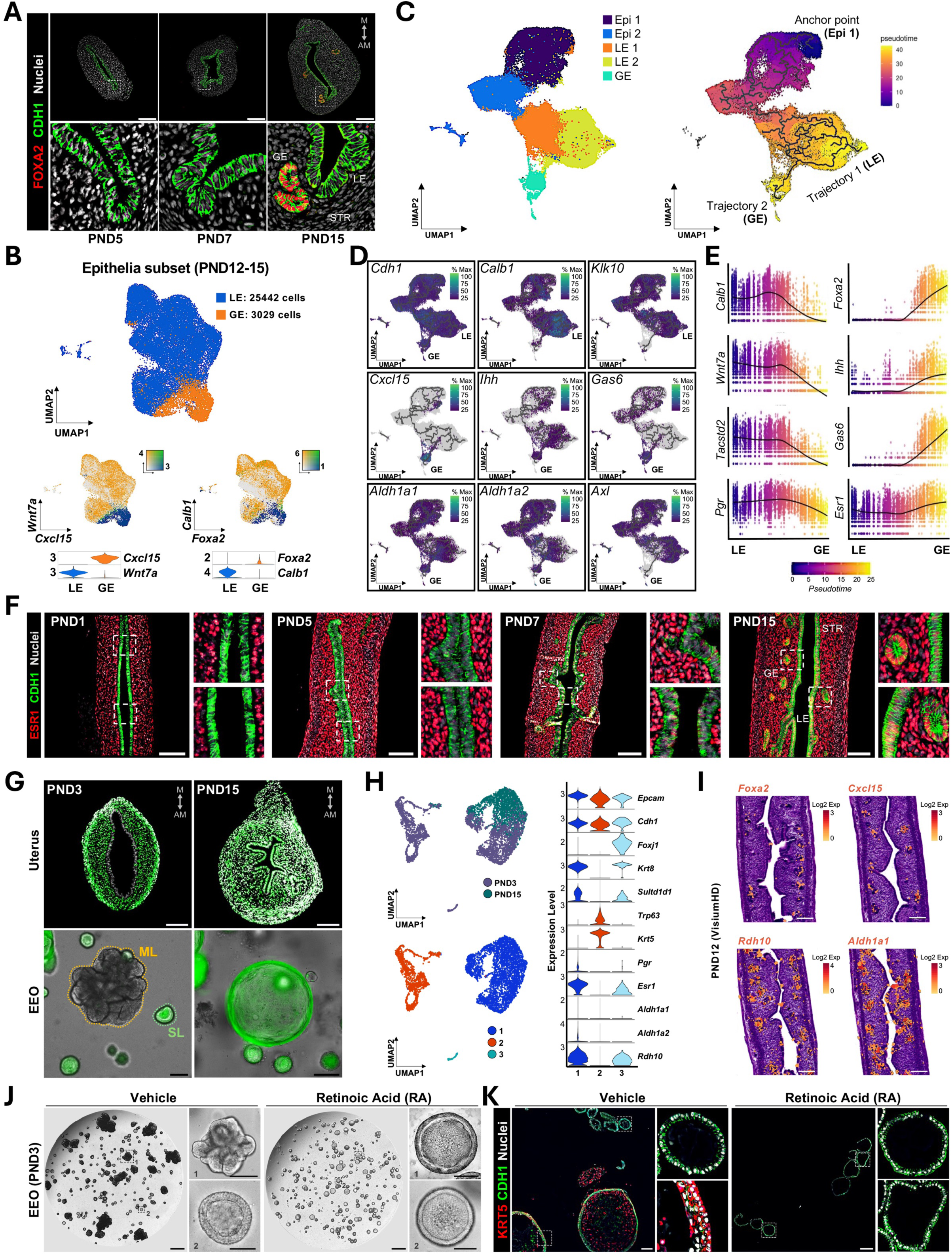
ESR1 acquisition and retinoic acid signalling suppress a developmentally transient epithelial state. (A) Representative immunofluorescence images of uterine sections at PND5, PND7, and PND15 showing FOXA2-positive glandular epithelium. Scale bars, 100 μM. (B) Uniform Manifold Approximation and Projection (UMAP) embedding of epithelial cells (n = 28,471) from PND12-D15 showing luminal epithelial (LE) and glandular epithelial (GE) populations. Feature and violin plots below depict lineage-specific markers. (C) Pseudotime analysis of epithelial differentiation coloured by subepithelial type (left n = 63,902) or illustrating trajectory modelling (right) from a common epithelial progenitor pool toward luminal (LE; Trajectory 1) and glandular (GE; Trajectory 2) fates. (D-E) Feature plots showing expression of selected genes along the luminal and glandular trajectories (D) and expression dynamics of selected luminal and glandular genes markers across pseudotime (E). (F) Representative immunofluorescence images of uterine sections from PND1 to PND15 showing progressive epithelial acquisition of ESR1. Scale bars, 100 μM. (G) Representative images of uteri (top; scale bars, 100 μM) and endometrial epithelial organoids (bottom; scale bars, 100 μM) derived from *Esr1^Cre^;Sun1-GFP* mice at PND3 and PND15. (H) scRNA-seq reanalysis of PND3 and PND15 EEO from GSE278635^22^. UMAP embedding of epithelial cells (n = 5764) coloured by developmental timepoint (top) or subcluster identity (bottom), and violin plots showing expression of selected genes (right). (I) Visium HD spatial transcriptomic analysis of PND12 uteri. Note the gland-enriched expression of *Foxa2*, *Cxcl15*, *Rdh10*, and *Aldh1a1*. Scale bars, 100 μM (J) Representative brightfield images of PND3-derived EEO treated with vehicle or retinoic acid (RA). Scale bars, 500 μM; insets, 100 μM. (K) Immunofluorescence localization of KRT5 in PND3-derived EEO treated with vehicle or retinoic acid (RA). Scale bars, 100 μM Abbreviations: EPI, epithelium; GE, glandular epithelium; LE, luminal epithelium; STR, stroma; EEO, endometrial epithelial organoid; ML, multilayered; SL, single-layered; RA, retinoic acid, M, mesometrial; AM, anti-mesometrial.

Stromal cell subclustering similarly revealed a temporal transition from unorganized mesenchyme to a mature differentiated stromal compartment (Fig. 2B). Three early mesenchymal clusters (Mes_1-3), enriched from PND1 through PND5, expressed *Pdgfra*, *Wnt5a*, *Hoxa11,* and *Top2a* alongside transient markers that include *Efnb2*, *Pakap*, and *Frem1*, consistent with a proliferative, undifferentiated mesenchyme identity (Fig. 2B). The three stromal clusters (Str_1-3) expanded progressively from PND5 to PND15 and were defined by high expression of *Col1a1*, *Dcn*, *Lum*, and *Wnt16* with corresponding GSEA enrichment for Extracellular Matrix Organization, Vasculature Development, and Cell Migration, reflecting acquisition of a mature stromal identity capable of providing structural and paracrine support to the overlying epithelium^13^ (Fig. 2B and Sup. Fig. 3B-C). Epithelial-Mesenchymal Transition and Cholesterol Homeostasis hallmarks were enriched across timepoints, suggesting these programs are sustained across the mesenchymal to mature/patterned stromal trajectory (Sup. Fig. 3B-C).

### Epithelial-Mesenchymal Crosstalk Governing Uterus Development

CellChat^50^ analysis of ligand-receptor interactions between epithelial and mesenchymal compartments was performed to characterize intercellular crosstalk underlying uterine development. Stromal clusters dominated outgoing communication at E16.5 through PND5, consistent with mesenchyme-driven instruction of the early epithelium, while bidirectional epithelial-stromal communication emerge progressively by PND12-15 (Sup. Fig. 3D–F). This shift suggests that epithelial cells transition from passive to active participants in the developmental signaling underlying the specification of luminal and glandular fates. Changes in TGFB, canonical and non-canonical WNT, BMP, GDF, IGF, FGF, GAS, and PROS signaling pathways were observed across timepoints (Sup. Fig. 3F). At PND5, mesenchymal *Tgfb2* and *Tgfbi* signaled through epithelial *Tgfbr2* and *Tgfbr3*, implicating stromal TGFB as an early mesenchymal-to-epithelial patterning cue, as well as TWEAK signaling via *Tnfsf12a-Tnfrsf12a*, consistent with active stromal remodeling during early neonatal organogenesis^13^ (Fig. 2C–D). As the epithelium matured and glands emerged, HGF signaling through *Hgf*-*Met* was found to be prominent in the mesenchyme-to-epithelium axis, while Hedgehog (HH) signaling appeared exclusively from the GE to the stroma via *Ihh*-*Ptch1* (Fig. 2E–F), recapitulating the canonical IHH-PTCH1 axis well characterized in the adult uterus^51–53^. Notably, GAS signaling via the *Gas6-Axl* axis underwent a fundamental shift from *Gas6* outgoing from the mesenchyme at PND5 to GE-derived signaling acting on both the luminal epithelium and stroma by PND12-15 (Fig. 2C–F). This reorganization was further reflected in the cross-species comparison, where *Gas6* was among the least conserved genes (Sup. Fig. 2E), consistent with its post-adenogenesis shift from stromal to glandular origin representing a developmental transition not yet captured in human fetal uterine samples at PCW 6-20 (Fig. 1F-H).

Visium HD spatial transcriptomics (10X Genomics) was performed on longitudinal sections of PND5 and PND12 uteri to add spatial context to this predicted cellular crosstalk (Fig. 2G–H). At PND5, *Gas6* transcripts were detected diffusely within the mesenchyme, while by PND12 *Gas6* was detected in the GE and stroma. Similarly, *Ihh* expression was within the GE while *Ptch1* was detected in the adjacent stroma, consistent with the emergence of GE-to-stroma IHH-PTCH1 signaling during this developmental window (Fig. 2H). Together, these findings suggest that stromal-epithelial cell crosstalk undergoes directional changes during postnatal uterine development, with the GE progressively acquiring an active signaling role coincident with adenogenesis^11,16,17,27,49^, raising the question of how LE cell progenitors become transcriptionally specified toward a GE cell fate (Figs. 3A-B; Dataset 2).

### Transcriptional Divergence in Luminal and Glandular Epithelial Cell Fates

Epithelial cells were subjected to pseudotime analysis using Monocle3^54^ to define the transcriptional programs underlying LE and GE differentiation (Fig. 3C). Two trajectories expressing the LE marker *Cdh1* arose from a common progenitor pool, either continuing down the luminal (Trajectory 1) or progressing toward a glandular (Trajectory 2) cell fate (Fig. 3C-D). Along the LE trajectory, *Calb1*, *Klk10*, *Wnt7a*, and *Tacstd2* were upregulated, while the GE trajectory was defined by the acquisition of *Foxa2*, *Cxcl15, Aldh1a1, Aldh1a2*, *Gas6*, and *Ihh* (Fig. 3D-E). Notably, *Esr1* was enriched along the GE pseudotime trajectory while *Pgr* was progressively downregulated (Fig. 3E). These analyses suggest that GE lineage bifurcation involves a coordinated transcriptional program distinct from that underlying commitment to the LE lineage, consistent with previous work from our group and others, implicating estrogen and retinoic acid (RA) signaling as candidate regulators of GE cell fate maintenance and uterine epithelial identity^11,16,23,34,55,56^

### ESR1 Expression During Epithelial Differentiation

The emergence of *Esr1* along the GE pseudotime trajectory prompted examination of ESR1 expression during postnatal development (Fig. 3F). At PND5, ESR1 protein was not detectable in the uterine epithelium but was present in the surrounding mesenchyme, consistent with previous observations [for review^16,57,58^] and scRNA-seq indicating low *Esr1* transcripts in epithelia from early timepoints (Fig. 2A and Sup. Fig. 4A). ESR1 was first detected in the epithelium on PND7, coinciding with the emergence of nascent GE buds^59^, and was broadly expressed in all major uterine cell types by PND15 (Fig. 3F). This progressive epithelial acquisition of ESR1 is particularly significant in light of prior findings *in vitro* demonstrating that the capacity of the uterine epithelium to form multilayered organoids, which reflects a transient epithelial cell state characterized by luminal-to-basal differentiation, is high in organoids derived from PND5 uteri but declines sharply by PND7 and is virtually absent by PND15^22^. The inverse relationship between epithelial ESR1 expression onset and multilayered EEO-forming capacity across this developmental window suggests a potential role for ESR1 in the progressive loss of epithelial plasticity described in organoids from the newborn mouse uterus^22^. In line with this observation and consistent with our previous work^23^, organoids established from *Esr1*-null mice at PND15 retained multilayered-forming capacity beyond the normal neonatal window. Interestingly, *Pgr*-null organoids exhibited a single-layered phenotype indistinguishable from wild-type controls (Sup. Fig. 4B), indicating that ESR1, but not PGR, is required to restrict this transient cell state. Notably, p63 protein was not detected within the uterine epithelium at any postnatal timepoint examined but was present in the cervical epithelium, suggesting that the transient cell population expanded in neonatal organoids does not correspond to a spatially discrete p63-positive compartment within the uterus proper *in vivo* (Sup. Fig. 4D–E).

**Figure 4.**
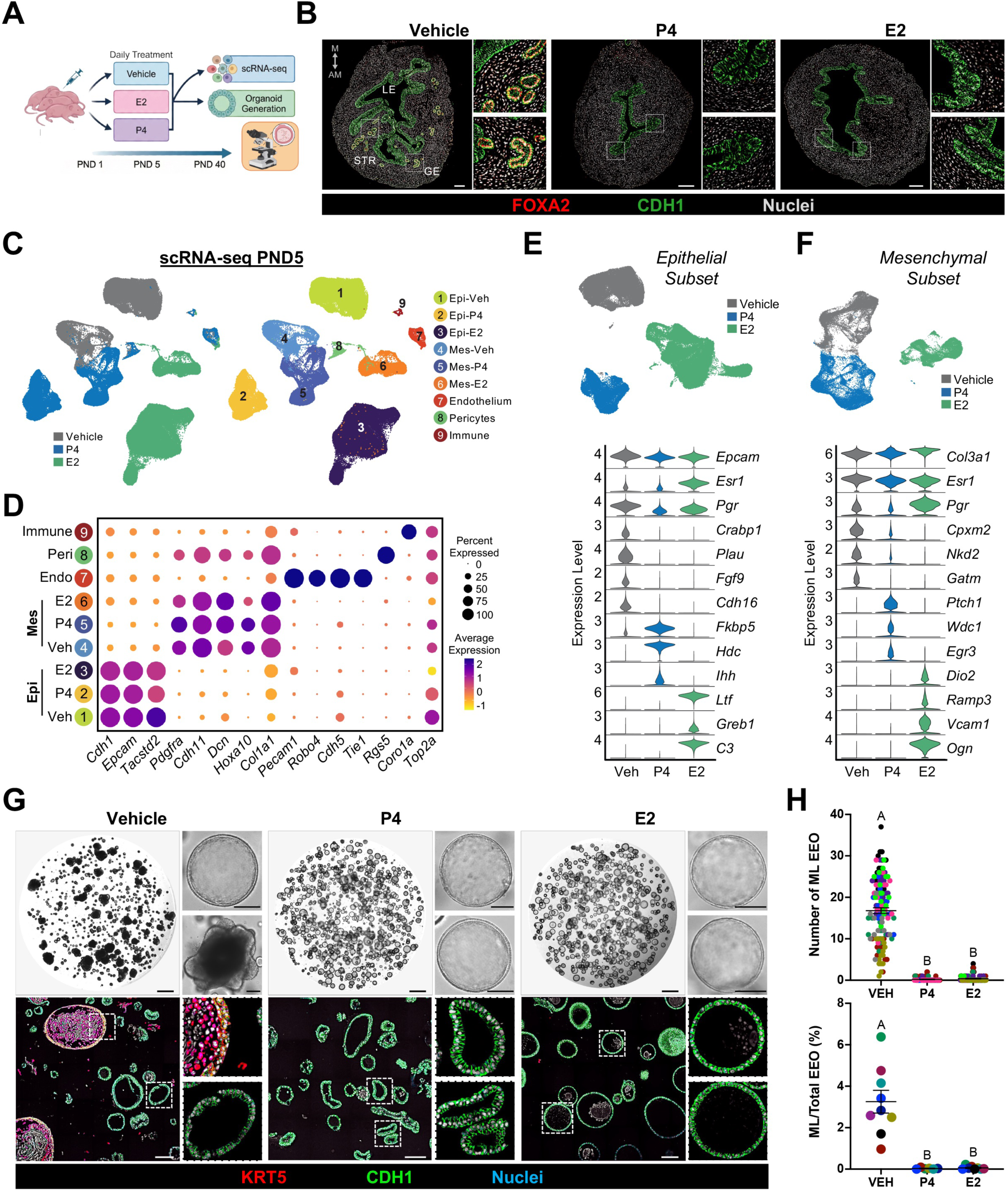
Neonatal steroid hormone exposure disrupts adenogenesis and abolishes the multilayered organoid phenotype. (A) Schematic of the neonatal steroid hormone treatment. Female mice were treated daily from PND1 to PND5 with vehicle, progesterone (P4), or estradiol (E2), followed by tissue collection for histological, transcriptomic, and organoid analyses. (B) Immunofluorescence localization of FOXA2 in PND40 uterine sections following neonatal P4 or E2 exposure relative to vehicle-treated controls. Scale bars, 100 μM. (C) Uniform Manifold Approximation and Projection (UMAP) embedding of PND5 uterine cells (n = 144,808) after neonatal vehicle, P4, or E2 treatment, coloured by treatment condition (left) or cell cluster identity (right). (D) Dot plot showing scaled average expression (color) and percentage of cells expressing (dot size) selected marker genes across epithelial and mesenchymal subsets. (E-F) Top, UMAP embedding of epithelial cells (E; n = 95,499) and mesenchymal cells (F; n = 44,371) coloured by treatment. Bottom, violin plots of selected epithelial (E) and mesenchymal (F) genes showing hormone-specific transcriptional responses. (G) Representative brightfield (top) and immunofluorescence (bottom) images of EEO established from vehicle, P4, or E2 treated mice at PND5. Vehicle-treated controls produced both single-layered and KRT5-positive multilayered EEO, whereas organoids from P4 and E2 treated mice lacked ML structures. Scale bars, 500 μM; insets, 100 μM. (H) Top, quantification of multilayered EEO number; each dot represents a EEO drop or technical replicate (n = 8-20 EEO drops per biological replicate per treatment). Bottom, percentage of multilayered EEO; solid dots represent biological replicates (n = 8-20 technical replicates per biological replicate per treatment). All experiments were performed on passage 3 organoids (n = 6-9 biological replicates per treatment). Data are presented as mean ± SEM. Letters indicate statistical differences between groups (*p* < 0.05; ANOVA with the Bonferroni multiple-comparison test). Abbreviations: EEO, endometrial epithelial organoid; ML, multilayered; EPI, epithelium; MES, mesenchyme; VEH, vehicle; P4, progesterone; E2, estrogen; M, mesometrial; AM, anti-mesometrial.

To determine the lineage origin of the plastic progenitor population, endometrial epithelial organoids (EEO) were derived from uteri of *Esr1*^Cre^;Sun1^GFP^ mice on PND3 and PND15 (Fig. 3G). Organoids from PND3 mice formed both single layered and multilayered structures. Multilayered EEO were GFP-negative, indicating they developed from ESR1 lineage-negative cells, while single layered organoids were GFP-positive. In contrast, all organoids were single layered and GFP-positive from PND15 uteri, revealing that ESR1 lineage-negative cells capable of multilayered EEO formation are lost as epithelial ESR1 expression is acquired across development (Fig. 3F-G). Furthermore, reanalysis of a scRNA-seq dataset from neonatal EEO (GSE278635)^22^ confirmed that PND3 cultures contained a transcriptionally distinct population expressing the basal markers *Trp63* and *Krt5* that lacked *Esr1*, *Pgr*, and RA metabolism genes including *Aldh1a1*, *Aldh1a2*, and *Rdh10*, whereas PND15 organoids lacked the basal gene signature (Fig. 3H; Sup. Fig. 4C). These results support the idea that multilayered EEO arise from a developmentally transient ESR1 lineage-negative epithelial cell population found only in the uterus during the first week of postnatal life.

### Retinoic Acid Signaling Suppresses Multilayered Organoid Formation

The absence of RA metabolism genes in multilayered EEO contrasted with their enrichment along the GE pseudotime trajectory (Fig. 3D) and their spatial restriction to the GE on PND12 (Fig. 3I), implicating RA signaling as a potential driver of epithelial lineage specification. To test this functionally, EEO established from PND3 uteri were treated with all-trans-retinoic acid (atRA). In contrast to the multilayered organoids developed in control cultures, only single layered organoids were found in cultures treated with atRA, phenocopying the morphology of EEO derived from PND15 uteri^22^ (Fig. 3J). Treatment with atRA resulted in the absence of KRT5-positive cells in PND3 organoids, suggesting that RA signaling is sufficient to suppress multilayered organoid-forming capacity (Fig. 3J-K). These findings are consistent with previous studies indicating a role of RA and estrogen signaling in the maintenance of epithelial homeostasis in the adult uterus^34^. Together, these results validate the neonatal organoid system as a functional screening platform for regulators of uterine epithelial cell fate identified by scRNA-seq. Acquisition of ESR1 expression and RA signaling emerge as two independent pathways that suppress a developmentally transient cell state in the epithelium as it becomes specified and lineage restricted in the developing neonatal uterus (Figs. 3F-K).

### Effects of Neonatal Steroid Hormone Exposure on Adenogenesis and Organoid Differentiation

Building on the finding that multilayered EEO arise from a developmentally transient ESR1-negative epithelial cell state that precedes GE lineage bifurcation^11,16,17,27,49^, we determined if hormone disruption of adenogenesis *in vivo* produces a corresponding loss of multilayered organoid-forming capacity, which would functionally connect this *in vitro* cell state to the *in vivo* GE specification program. To test this, female mice were administered daily subcutaneous injections of progesterone (P4; 50 μg/g body weight) or 17-beta-estradiol (E2; 8 μg/g body weight) from PND1 through PND5, the developmental window during which the ESR1-negative progenitor population is active (Fig. 3 G-K) and formation of multilayered EEO is highest^22^. Neonatal P4 exposure inhibits adenogenesis in the uterus of both sheep and mice^16,60–63^. A marked reduction in FOXA2-positive GE structures were observed in mice treated with either P4 or E2 relative to vehicle controls on PND40, confirming that the five-days neonatal steroid hormone treatment effectively disrupts the GE lineage specification program *in vivo* (Figs. 4A-B).

To characterize the transcriptional basis of this disruption, scRNA-seq was performed on uteri collected at PND5 following the treatment period (Fig. 4C-D) using an EpCAM-based magnetic sorting approach (Fig. 1A). After quality control and filtering (see Methods), a total of 144,808 cells were retained for downstream analysis. While no major differences were observed in endothelial, perivascular, and immune cells, unsupervised clustering revealed that P4 and E2 produce fundamentally distinct transcriptional states in both epithelial and mesenchymal compartments (Fig. 4C-F; Dataset 3). Within the epithelial cell subset (n=95,499 cells), genes that define uterine neonatal epithelial identity in normal control uteri, including *Crabp1*, *Plau*, *Fgf9*, and *Cdh16,* were lost in both P4 and E2 treatment groups, suggesting a common suppression or elimination of the normal pre-adenogenesis epithelial state (Figs. 2A, 3C-E, and 4E). Treatment with P4 induced *Fkbp5*, *Hdc,* and *Ihh,* and E2 upregulated classical E2-responssive genes including *Ltf*, *Greb1*, and *C3*, as well as *Esr1* (Fig. 4E). Further, ESR1 protein was observed in the uterine epithelium of E2-treated mice, a timepoint at which ESR1 is normally restricted to the stroma (Fig. 3F, Fig. 4E-F, and Sup. Fig. 5A). In the mesenchymal clusters (n=44,371 cells), E2 increased *Ogn*, *Dio2*, and *Ramp3* together with *Pgr*, consistent with E2 effects on PGR in the stroma of adult mouse uteri^64,65^. The E2 increase in *Pgr* mRNA was validated by IF (Fig 4F, Sup. Fig. 5A). Treatment with P4 upregulated *Ptch1*, *Wdc1*, and *Egr3* expression in the mesenchyme (Fig. 4F). This premature epithelial *Ihh* to mesenchymal *Ptch1* axis at PND5 recapitulates in a disorganized fashion the signaling architecture that normally emerges exclusively from committed GE cells after PND12 (Fig. 2C-H, Fig. 3C-E, and Fig. 4E), suggesting that exogenous P4 short-circuits the normal developmental sequence of GE specification. This single-cell resource is also accessible at www.genesearch.org/sklab/glanddisruption/.

**Figure 5.**
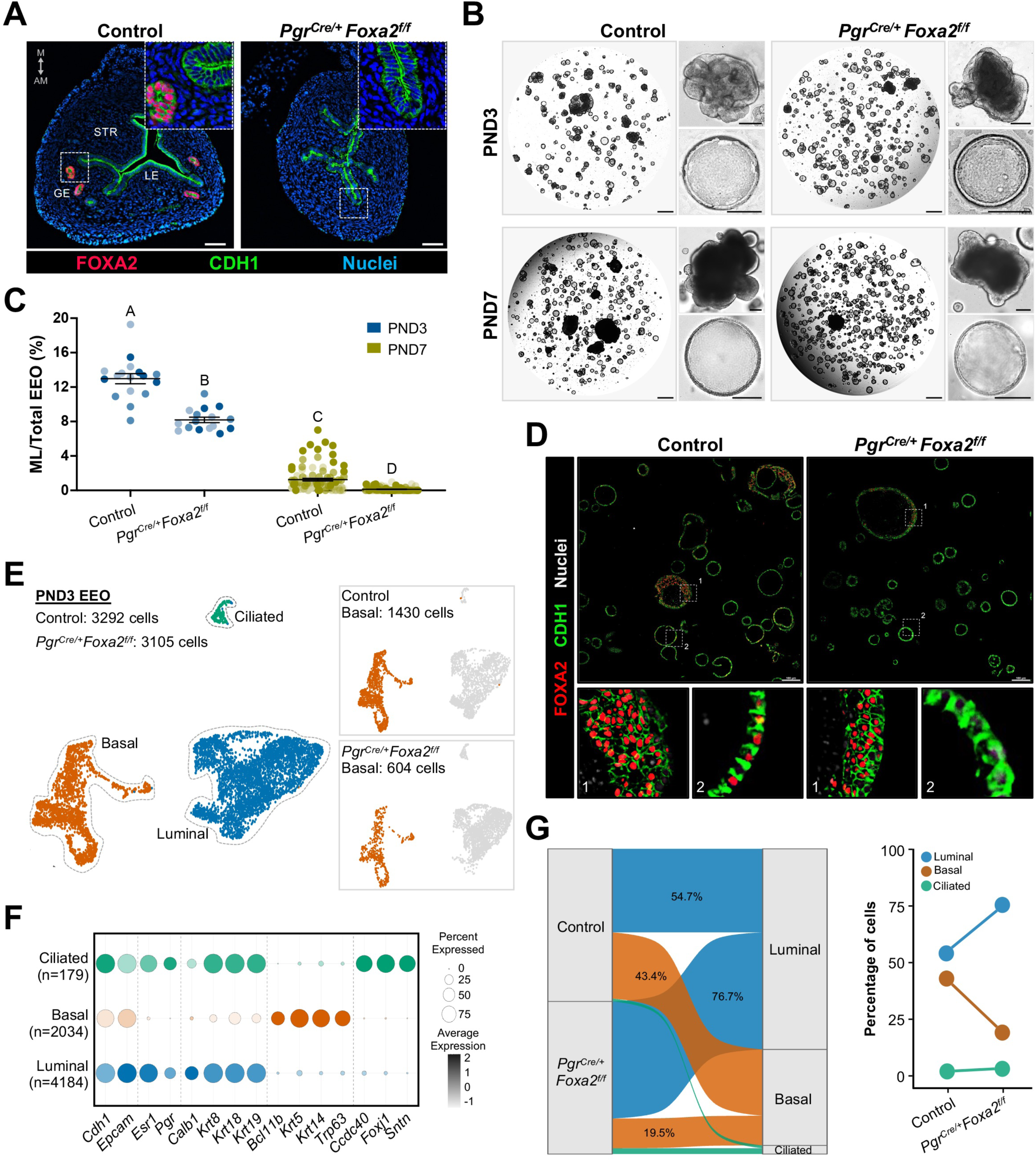
Genetic Ablation of Glandular Epithelium Impacts Neonatal EEO Phenotype. (A) Immunofluorescence localization of FOXA2 in wild-type control and *Pgr^Cre/+^Foxa2^f/f^* uterine sections at PND15. Scale bars, 50 μM. (B) Representative brightfield images of PND3 (top) and PND7 (bottom) EEO established from control (left) or *Pgr^Cre/+^Foxa2^f/f^* (right) mice. Scale bars, 500 μM; insets, 100 μM. (C) Percentage of multilayered EEO formation in cultures established from control and *Pgr^Cre/+^Foxa2^f/f^* mice at PND5 (blue) and PND7 (green). Each dot represents an EEO drop or technical replicate (n = 4-20 EEO drops per biological replicate per genotype), and color shading indicates biological replicates. All experiments were performed on passage 3 organoids (n = 4-8 biological replicates per genotype per PND). Data are presented as mean ± SEM. Letters indicate statistical differences between groups (*p* < 0.05; ANOVA with the Bonferroni multiple-comparison test). (D) Immunofluorescence localization of FOXA2 in control and *Pgr^Cre/+^Foxa2^f/f^* PND7 EEO. Scale bars, 100 μM. (E-F) Uniform Manifold Approximation and Projection (UMAP) embedding of epithelial cells (E, n = 3105) from PND3 EEO established from control and *Pgr^Cre/+^Foxa2^f/f^*mice and violin plots (F) of selected genes indicating luminal, basal, and ciliated cells clusters. (G) Alluvial plot (left) and quantification (right) showing the proportions of luminal, basal, and ciliated cells in EEO derived from Control and *Pgr^Cre/+^Foxa2^f/f^* mice at PND3, demonstrating a shift toward luminal identity and a reduction in basal cell frequency upon *Foxa2* loss. Abbreviations: LE, luminal epithelium; GE, glandular epithelium; STR, stroma; EEO, endometrial epithelial organoid; M, mesometrial; AM, anti-mesometrial.

CellChat^50^ analysis also revealed that hormone exposure disrupts epithelial and mesenchymal cell communication in the neonatal uterus. In vehicle-treated control uteri, outgoing epithelial signaling was defined by KLK, PLAU, WNT, GDF, and EGF pathways (Sup. Figs. 5B-C). These were identified as determined LE markers in the developmental atlas and pseudotime (Fig. 2A; Fig. 3D–E), establishing them as core communication signals of the pre-adenogenesis uterine epithelium, and both P4 and E2 converged on disruption of this core signaling identity despite their distinct transcriptional mechanisms^57,66^ (Sup. Fig. 5B-C). Further, P4 and E2 induced distinct aberrant signaling programs. Stromal outgoing communication was largely preserved under P4, suggesting that P4 acts predominantly through epithelial transcriptional reprogramming rather than by disrupting stromal instructive signals (Sup. Fig. 5B-C). Conversely, E2 induced a wider disruptive reorganization resulting in aberrant inflammatory signaling through IFN-I and TRAIL in the epithelium while simultaneously rewiring stromal GAS, PTN, and SLIT instructive pathways (Sup. Fig. 5B-C), which normally coordinate mesenchymal-to-epithelial cell communication during this developmental window (Sup. Fig. 3D-F). Together, these data confirm that while P4 and E2 produce molecularly distinct perturbations of the neonatal uterine signaling landscape, both converge on the elimination of the normal pre-adenogenesis epithelial identity, pushing the epithelium into aberrant transcriptional states during critical windows of specification in the uterus. (Fig. 4 E-F; Sup. Fig. 5A-C).

To determine whether these *in vivo* molecular perturbations translate to loss of the developmentally plastic epithelial state evident by multilayered EEO formation *in vitro,* EEO were established from vehicle, P4, and E2-treated mice at PND5. All three groups formed normal single-layered cystic structures, confirming that neonatal hormone exposure does not impair overall organoid formation (Fig. 4G). However, while vehicle-treated controls produced both single-layered and KRT5-positive multilayered EEO, consistent with activation of a luminal-to-basal differentiation program (Fig. 4G), organoids from mice treated with P4 and E2 were devoid of multilayered structures, resembling those derived from PND15 uteri (Fig 3G, Fig. 4G-H). This selective loss of multilayered organoid-forming capacity was shared between treatment groups despite their distinct *in vivo* transcriptional signatures, demonstrating that the multilayered EEO phenotype is functionally associated with the GE specification program and is lost as a common consequence of disrupting adenogenesis, regardless of the specific hormonal perturbation employed (Fig. 4E-H).

### Genetic Ablation of Glandular Epithelium Impacts Neonatal EEO Phenotype

The consistent loss for the capacity of multilayered organoids to develop from neonatal uteri observed alongside disruption of glandular morphogenesis prompted us to ask whether this functional relationship is governed by a shared transcriptional mechanism. FOXA2 is a transcription factor required for uterine gland formation^27,67,68^ and its expression defines the GE pseudotime trajectory (Fig. 3C-E), making it a compelling candidate to test whether the multilayered EEO state is associated to the GE specification program. EEO were established from the uteri of *Pgr^Cre^;Foxa2^f/f^*conditional knockout mice on PND3 and PND7. In this mouse model, FOXA2 is deleted during the peri-adenogenesis window and results in failure of uterine gland differentiation^15^. Note the absence of FOXA2-positive glands in the uterus of PND15 *Pgr^Cre^;Foxa2^f/f^*mice (Fig. 5A).

Uterine epithelial cells from both PND3 and PND7 *Pgr^Cre^;Foxa2^f/f^*mice formed lower (P<0.05) numbers of multilayered organoids relative to the vehicle controls (Figs. 5B-C). This phenocopies the loss of multilayered organoid-forming capacity observed following neonatal P4 and E2 treatment (Fig. 4G-H), establishing that the coupling between GE specification and multilayered organoid-forming capacity holds across both hormonal and genetic models of disrupted adenogenesis. Immunofluorescence on EEO derived from *Pgr^Cre^;Foxa2^f/f^*mice revealed that the residual multilayered structures retained FOXA2 expression while single-layered organoids were FOXA2-negative, contrasting with control cultures where FOXA2 is present in multilayered and a subset of single layered EEO (Fig. 5D). As the basal cell state identified by scRNA-seq lacks PGR expression and would therefore not be targeted by *PgrCre*-mediated recombination (Fig. 3H; Sup. Fig. 4C), the residual multilayered EEO in this model are consistent with cells that do not express *Pgr* at the time of uterine isolation^69^. This finding supports the idea that multilayered organoid-forming capacity is intrinsic to a *Pgr*-negative, yet FOXA2-competent transient epithelial cell state (Fig. 5D).

Next, scRNA-seq was performed on organoids derived from PND3 control and *Foxa2* conditional knockout mice. Three transcriptionally distinct epithelial populations were identified across both groups: a luminal cluster (4,184 cells; *Cdh1*, *Epcam*, *Calb1*), a basal cluster (2,034 cells; *Krt5*, *Krt14*, *Trp63*, *Bcl11b*), and a ciliated cluster (179 cells; *Foxj1*, *Sntn*) (Figs. 5E-F; Dataset 4). The proportion of cells in the basal cluster was lower in EEO from uteri of *Foxa2* conditional-deletion mice than controls (43.4% vs 19.5%), whereas the luminal cells were higher in *Foxa2* conditional knockout mice (54.7% vs. 76.7%) (Fig. 5E-G). These shifts in organoid composition parallel the reduction in multilayered organoid formation observed morphologically (Fig. 5B-C). Thus, genetic ablation of GE differentiation may selectively deplete cells that possess a basal cell identity or developmental program with multilayered organoid-forming ability. Taken together, these results suggest independently through neonatal steroid hormone exposure and conditional genetic deletion of *Foxa2* that the capacity for multilayered organoids to arise from neonatal uteri is linked to the GE lineage specification program. Combined with the developmental cell atlas, pseudotime trajectory analysis, lineage tracing, and RA signaling data presented above, these findings identify the multilayered EEO phenotype as the *in vitro* correlate of a developmentally transient epithelial cell state that precedes and is required for glandular fate specification in the postnatal uterus.

## DISCUSSION

The specification of glandular epithelium from luminal progenitors during uterine development is essential for women’s health and fertility, yet the transcriptional trajectories, epithelial-mesenchymal signaling dynamics, and transitional cell states that govern this process from fetal development to adulthood remain limited^11–13^. This gap is particularly consequential in humans where uterine adenogenesis initiates during the third trimester of pregnancy, a window that is clinically vulnerable to endocrine disruption and inaccessible to functional study^45–47,70,71^. Here, a single-cell roadmap of mouse uterine development from undifferentiated Müllerian Duct through completed postnatal uterine patterning^1^ was generated. Characterization of the progressive differentiation of the epithelial and mesenchymal compartments provides new insights into epithelial-mesenchymal communication that governs uterine development and dynamically changes during GE lineage bifurcation. By integrating organoid culture, lineage tracing, and hormonal and genetic perturbations, the studies reveal a transient ESR1-negative epithelial cell state that is present during the early pre-adenogenic period and progressively lost as uterine development proceeds. Further, the data demonstrate that epithelial ESR1 acquisition and RA signaling each suppress epithelial plasticity and promote determination and maturation to uterine type luminal epithelium. Cross-species comparison with a recently published atlas of fetal human FRT^14^ reveals conservation of core epithelial and mesenchymal programs, contextualizing these findings within human uterine biology and identifying the pre-adenogenesis window as a critical yet largely undercharacterized determinant of GE development.

The progressive acquisition of ESR1 in the uterine epithelium during neonatal development coincides with loss of multilayered EEO-forming capacity, implicating ESR1 as a regulator that restricts epithelial plasticity and promotes epithelial determination and lineage bifurcation. This interpretation is supported by the observation that ESR1 loss in adult mice results in formation of multilayered organoids beyond the normal developmental window and is consistent with established roles of ESR1 in maintaining epithelial homeostasis in the adult uterus^23,55,56,72,73^. Independently, RA signaling was decreased in multilayered EEO *in vitro* and enriched in the GE *in vivo*. Consistent with these observations, addition of atRA to organoid cultures suppressed the transient epithelial state in neonatal EEO cultures, supporting a role for RA signaling in promoting epithelial maturation early in development^74,75^. The finding that ESR1 acquisition and RA signaling suppress multilayered EEO formation suggests that multiple developmental inputs act to limit this transient cell state to the first week of postnatal uterine development^22^. The transient cell state lacks both *Esr1* and *Pgr* expression, suggesting that it exists in the undetermined epithelium of the newborn uterus before GE differentiation. This is further supported by ovariectomy studies in newborn mice finding that FRT morphogenesis and fate specification in the neonatal uterus occur independently of ovarian steroid hormones^16,76^.

Tissue recombination studies established that the mouse FRT epithelium loses competence to respond to heterotypic mesenchyme induction of cell fate after PND7^18–21^. This loss of competence is mirrored by the age-dependent decline in formation of neonatal multilayered EEO^22^, yet the mechanisms restricting this developmental window of epithelial plasticity within the uterus remain unclear^24–26^. The present study identifies epithelial acquisition of ESR1 as an intrinsic signal that suppresses uterine epithelium plasticity. The cell state captured by neonatal EEO exhibits luminal-to-basal differentiation marked by KRT5 and p63 expression and is not restricted to early uterine development. Multilayered EEO with this signature also arise from adult mice with *Tgfbr2*-mediated endometrial hyperplasia and in humans from patient-derived endometrial cancer organoids^22,36^. Moreover, loss of ESR1 has been linked to upregulation of basal invasive markers in breast cancer^77–79^ and women with *Esr1*-negative endometrioid endometrial cancer are diagnosed with higher grade and advanced disease stages^80^. This suggests that basal cell differentiation, latent during normal uterine development, is co-opted under pathological conditions, analogous to mechanisms underlying epithelial dysfunction in other organ systems^81–87^. While our understanding of the molecular basis governing this latent cell state in the endometrium remains limited, these observations position ESR1 as a conserved regulator of epithelial cell fate across development and disease. Together, these findings reframe the multilayered EEO phenotype as a functional readout of a biologically relevant developmental transition with implications for epithelial homeostasis, rather than an *in vitro* artifact, supporting foundational theories of FRT development^18–21^.

Epithelial-stromal signaling undergoes coordinated reorganization during maturation and bifurcation of epithelial cell lineages in the mouse uterus^11–13^. Our analysis extends prior genetic studies [for review^41,44,88–98^] by showing that, as LE and GE lineages emerge, the epithelium transitions from a largely passive recipient of stromal cues to an active signaling compartment. *Ihh-Ptch1*, a well-established mediator of P4 action in the adult uterus, emerges in the GE after PND12, whereas GAS signaling similarly shifts from stroma to glands as adenogenesis proceeds^51–53,65,99^. The progressive enrichment of *Gas6* in the GE raises the possibility that gland-derived GAS6 acts through AXL to support epithelial maintenance after lineage commitment. Neonatal hormone exposure disrupted both this epithelial-stromal crosstalk and the core transcriptional identity of the neonatal epithelium. Specifically, P4 induced premature *Ihh-Ptch1* signaling at PND5, a configuration normally observed only after glandular commitment. Because conditional ablation of *Ihh* confers resistance to P4-mediated suppression of uterine adenogenesis, premature activation of this pathway may contribute to failed glandular specification^100^. Together with the finding that both P4 and E2 eliminate multilayered EEO formation and disrupt GE development, these data support a model in which successful GE specification requires precise temporal coordination of cell-intrinsic transcriptional programs and epithelial-stromal communication.

Human uterine gland development initiates around 20-22 PCW and is completed only later in postnatal maturation^45–47,70,71^, and the conserved epithelial programs identified between the developing mouse and human fetal epithelium suggest that the transcriptional programs defined here are likely operative during an analogous human pre-adenogenic window^14^. The findings that neonatal exposure to exogenous P4 or E2 disrupts the transient epithelial cell state and GE morphogenesis raise the possibility that equivalent exposures during the human fetal pre-adenogenesis window may compromise glandular specification with long-term consequences for uterine function and fertility^2,10^. Notably, daughters of women exposed to diethylstilbestrol have an increased risk of uterine and vaginal epithelial abnormalities, providing precedent that estrogenic disruption during critical developmental windows can have lasting consequences for epithelial differentiation^5,8,101,102^. Fetal exposure to diethylstilbestrol induced epithelial ESR1 expression in xenografts from 14-PCW human specimens^9,71^. This finding is consistent with the mouse data showing that neonatal E2 treatment induces premature epithelial ESR1 expression at PND5 and disrupts the pre-adenogenic epithelial state^58,103,104^. Likewise, progesterone supplementation is prescribed during the second and third trimester of pregnancy, yet existing follow-up studies assessed neurodevelopmental outcomes in children followed only to early childhood and did not examine FRT organs^105–107^, leaving the long-term consequences of gestational progesterone exposure for GE development and epithelial function as an unaddressed question^107,108^. These results highlight the need to define the molecular basis of this latent epithelial plasticity and its implications for women’s health. Because direct functional perturbation of the human fetal uterus is not feasible, the neonatal mouse organoid system validated here provides a tractable platform for identifying regulators of glandular specification relevant to human reproductive development and disease.

Taken together, this study provides a single-cell framework spanning fetal to postnatal uterine development and identifies a transient epithelial cell state governed by ESR1 acquisition and RA metabolism that is functionally coupled to glandular morphogenesis across independent hormonal and genetic perturbation models. While direct *in vivo* lineage evidence linking this transient cell state to the glandular epithelium remains to be established, the convergence of transcriptional, lineage tracing and functional evidence across multiple experimental systems supports its biological significance. Future studies characterizing the chromatin accessibility landscape of this transient cell state and its equivalent in human uterine fetal endometrial organoids and tissues will be essential for translating these mechanistic insights into a more complete understanding of glandular dysfunction. The pre-adenogenesis window emerges from these studies as a critical and previously undercharacterized determinant of adult uterine function whose disruption has consequences extending from failed glandular morphogenesis to the broader landscape of uterine factor infertility and endometrial disease^30,109–111^.

## LIMITATIONS

Several limitations of the current studies should be acknowledged. The developmental atlas spans E16.5 through PND15, capturing the initiation and histological progression of uterine adenogenesis but not terminal glandular differentiation, hormone responsiveness, or the pubertal maturation required for full reproductive competence. The spatial transcriptomic data add important regional resolution but are derived from two postnatal timepoints and do not capture the full dynamics of signaling reorganization across the fetal-to-neonatal transition. Cross-species conservation is inferred from transcriptional similarity between human fetal and mouse fetal-neonatal datasets. Whether the transient ESR1-negative epithelial cell state identified here has a direct functional counterpart in the human fetal uterus, and whether human fetal or neonatal organoids recapitulate the multilayered EEO phenotype, remain important open questions.

## METHODS

### Animals

All animal procedures were approved by the Institutional Animal Care and Use Committee of the University of Missouri, Columbia and were conducted according to the NIH Guide for the Care and Use of Laboratory Animals. C57BL/6J mice were obtained from The Jackson Laboratory (JAX #000664). Floxed *Foxa2* mice^112^ (JAX #022620) were crossed with *Pgr^Cre^*to generate conditional knockout animals. *Pgr^Cre^* mice^69^ were provided by Dr. Francesco Demayo (National Institute of Environmental Health Sciences, Durham, North Carolina) and Dr. John Lydon (Baylor College of Medicine, Houston, Texas). *Esr1* null (*Esr1^-/-^*) mice^103^ were provided by Dr. Dennis Lubahn (University of Missouri, Columbia, Missouri). *Esr1^Cre^* mice (JAX #017911) were crossed with *Sun1*-*GFP* mice^113^ (JAX#030952) for *Esr1*-lineage tracing studies. *Esr1^Cre^* mice^114^ were kindly provided by Dr. David Anderson (California Institute of Technology, Pasadena, California). For fetal collections, observation of a vaginal plug was defined as embryonic day E0.5, developmental stage was confirmed by the Theiler staging criteria at dissection, and mice were sexed by evaluating presence of testis or well-defined Müllerian ducts. For neonatal collections, pregnant dams were monitored daily from 16:30 to 18:00, and the day of birth was considered postnatal day (PND) 0 for all studies. For steroid hormone treatments, beginning at PND1, C57BL/6J female pups were weighed and administered daily subcutaneous injections of E2 (8 μg/kg body weight; E8875; Sigma-Aldrich), P4 (50 μg/g body weight; P0130; Sigma-Aldrich), or oil (V/V) until PND5. Hormone dosages were selected based upon previous reports^16^.

### Mouse endometrial epithelial organoids (EEO)

Uterine horns were dissected from female reproductive tracts as described previously^22^. Uteri were digested in 1% trypsin (Sigma, T4799) in calcium and magnesium-free HBSS (Gibco, 14175-095) for 45 minutes at 4°C followed by 45 minutes at 37°C using an orbital shaker. The digestion was stopped with 1% soybean trypsin inhibitor (Gibco, 17075-029) in HBSS with 5 mM MgCl_2_ and 0.1 mg/mL DNAse-I (Roche, 10104159001). For PNDs 3, 5, 7 (n= 3-5 mice/biological replicate; n= 4-18 biological replicates per PND), 10 μL pipette tips were carefully pressed using small forceps to create a slightly smaller diameter than the uterus, and endometrial epithelial sheets were retrieved by gentle suction using a P10 pipette. For PND15 (n= 2-3 mice/biological replicate; n= 6 biological replicates), the digested uterine horns were gently squeezed with #5 forceps. The cell pellets were rinsed gently with Base Organoid Media (Table S1). After centrifugation (300 x *g* for 3 minutes) and washing, epithelium pellets were resuspended in 80% Cultrex (R&D Systems, 3445-005-01) and 20% organoid expansion media (Table S2), and plated in 15 μL drops; 4 drops/well in 24-well plates. For retinoic acid treatment, EEO cultures were supplemented with 5uM of all-trans retinoic acid (atRA, Sigma, R2625) or vehicle control (DMSO, V/V) for 20 days prior to analysis.

Organoids were passaged (1:3 ratio) every 7-10 days as previously described^115^. For passaging, Cultrex/organoid drops were detached from the cell culture plate by gentle pipetting and transferred into 1.5 mL Eppendorf tubes. After centrifugation (300 x *g* for 3 minutes), the supernatant was removed, replaced by 1.5 mL of base organoid media, and pipetted up and down to dissociate the pellet. Following an additional centrifugation step, the cell pellet was resuspended in Cultrex and plated in 12-well plates (15 μL drops; 4 drops per well). Cultrex drops were incubated at 37 °C for 15 min prior to adding 800 μL of organoid expansion media (Table S2) per well. Organoid cryopreservation was performed with freezing medium consisting of 10% DMSO in fetal bovine serum (FBS, Sigma, F0926) as described previously^115^. All experiments were performed using passage 3 cultures. Brightfield images were captured on a Leica DMi8 inverted microscope and Leica K8 camera using Leica Application Suite X (LAS X). EEO number (n= 4-17 drops/biological replicate) and frequency of multilayered structures [(# of ML EEO) ÷ (total # of EEO)] were analyzed using ImageJ (n= 4-18 biological replicates/PND/phenotype/treatment) and the data are presented as the mean ± SEM. Statistical differences between groups were determined using one-way ANOVA with Bonferroni multiple-comparison test’s multiple comparisons test (GraphPad Prism 9) and statistical significance was defined as p < 0.05.

### Tissue digestion, single cell isolation, and sample processing for scRNA-seq

For scRNA-seq atlas studies, tissues from fetal (E16.5) and neonatal PND1 (n=90-125 uteri), PND5 (n=85-100 uteri), PND12 (n=45-60 uteri), and PND15 (n=36-45 uteri) C57BL/6J mice were digested using Liberase (0.2 mg/mL; Roche, 643505) diluted in phenol red-free RPMI 1640 medium supplemented with DNAse-I (0.5 mg/mL; Roche, 10104159001) for 45 min at 4°C, followed by 45 min at 37 °C using an orbital shaker (50 RPM). For exogenous E2 and P4 studies, hormone-treated C57BL/6J mice were collected 3 hours after the last injection on PND5 (E2, n=40-55 uteri; P4, n=60-75 uteri). The goal was to sequence a 50:50 ratio of epithelial to non-epithelial cells for all timepoints and conditions. To achieve this, digested tissues were subjected to magnetic separation using CD236 (EpCAM) microbeads following manufacturer’s instructions (Miltenyi Biotec, 130105958). EpCAM-positive epithelial cells bound to the magnetic columns and the flow-through containing the unlabeled non-epithelial cells were transferred into 5 mL tubes. Cell viability and efficiency of dissociation were determined using a Cellometer K2 Fluorescent Cell Counter (Revvity, CMT-K2-MX-150). Viability threshold for epithelial cells was set at or above 65%, while non-epithelial cells consistently showed 92% viability or above. Fetal cells did not pass this quality checkpoint (particularly with E16.5 epithelial cells showing lower than 30% viability). Therefore, magnetic sorting was not performed and, instead, E16.5 uteri (n=44-55 embryos) were digested and directly processed for sequencing as described below.

For each sample, 250,000 cells were fixed for 22 h at 4 °C, quenched and stored at −80 °C according to 10X genomic Fixation of Cells & Nuclei for Chromium Fixed RNA profiling (CG000478, 10X Genomics, Pleasanton, CA) using the Chromium Next GEM Single Cell Fixed RNA Sample preparation kit (PN-1000414, 10X Genomics). A total of ∼200,000 cells per sample were used for probe hybridization with mouse WTA probes (PN-1000496, 10X Genomics), pooled and washed following the Chromium Fixed RNA Profiling Reagent kit protocol (CG000527, 10X Genomics). GEMs were generated using Next GEM ChipQ (PN-1000422, 10X Genomics) on the Chromium X (10X Genomics) system with a target of at least 20,000 (10,000 epithelial: 10,000 non-epithelial) cells, according to the manufacturer instructions (CG000527, 10X Genomics).

### Organoid single-cell isolation and scRNA-seq

EEOs established from PND3 wild-type or *Pgr^Cre^Foxa2^f/f^*female mice were collected after 8 days of culture (n=10 females/PND) and processed for scRNA-seq as previously described^22^. Briefly, EEOs were enzymatically dissociated using 0.05% trypsin (40 min, 37°C), and centrifuged at 300 × g for 5 min. Cells were resuspended in Advanced DMEM/F12 medium supplemented with 1X antibiotic-antimycotic (Gibco, 15240062), passed through a 40 µm filter (Fisher, 087711) and pelleted (300 × g for 5 min). The cell pellets were then resuspended in 1 mL 0.04% BSA in PBS and counted to assess viability. Single-cell RNA sequencing was conducted at the University of Missouri Genomics Technology Core as described previously^22^. Briefly, single-cell droplet formation with a target count of 10,000 cells per sample was performed using the 10X Chromium system (10X Genomics; Pleasanton, CA). Sequencing was performed in paired-end mode using NovaSeq X Plus (Illumina).

### Visium HD spatial transcriptomics

Uterine horns from PND5 and PND12 females (*n* = 5 females per timepoint) were collected, fixed in 4% paraformaldehyde for 16 hours at 4°C, and processed for paraffin embedding. FFPE blocks were sectioned at 5 µm and mounted onto glass slides for histological evaluation or spatial transcriptomic profiling. Spatial Gene Expression library preparation and sequencing were performed by the Genome Technology Access Center at Washington University in St. Louis. Briefly, 5-µm paraffin sections were deparaffinized, stained with hematoxylin and eosin, and imaged to capture tissue morphology. Sections were processed using Visium HD H1 Slide, HD probe-based v1 chemistry, with the Visium Mouse Transcriptome Probe Set v2.0. Sequencing generated 159,452,943 reads for PND5 and 179,774,203 reads for PND12. Across the capture areas, 18,994 genes were detected in 208,076 8 µm binned squares under tissue for PND5, and 18,993 genes were detected in 293,387 8 µm binned squares under tissue for PND12.

### Bioinformatic analysis of scRNA-seq and 10x Visium HD data

For uteri scRNA-seq analysis, sample demultiplexing and read alignment were performed using the Cell Ranger (10X Genomics; v.9.0.1) multi-pipeline (cellranger multi) tool, which was configured via a sample-specific CSV configuration file to handle multiplexed library processing with the default parameters. Reads were aligned to the mouse reference genome (refdata-gex-mm10-2020-A), followed by filtering and counting of barcodes and unique molecular identifier (UMIs) to generate count matrices for each sample. For EEO scRNA-seq, the base call files were demultiplexed and processed to generate FASTQ files with Cell Ranger (v.9.0.1) command mkfastq, and then aligned to the to the mouse reference genome GRCm38 release 102 (Ensembl), followed by filtering and barcode plus UMI counting to generate count matrices for each sample. To remove ambient RNA contamination, the raw count matrices from Cell Ranger were processed using CellBender (v.0.3.2) with default parameters to generate denoised gene expression matrices. Processed count matrices were imported into R and analyzed using Seurat^116^ (v.5.3.0) for quality control and filtering using standard preprocessing workflow. Doublets were identified and removed using scDblFinder (v.1.20.2) with default parameters. Cells were retained based on the following quality control thresholds: number of detected genes between 200 and 15,000 (nFeature_RNA), total UMI counts between 0 and 100,000 (nCount_RNA), and mitochondrial read fraction below 10%. Doublets identified by scDblFinder were excluded prior to downstream analysis. Harmony^117^ was used for data integration when indicated and Loupe Browser files were created using LoupeR from 10X Genomics. Publicly available scRNA-seq datasets used in this study were downloaded from the NCBI database with accession codes E-MTAB-15475^14^, GSE278635^22^, and GSE275806^37^.

For epithelial sub-clustering, cells from postnatal timepoints (≥ PND1) were isolated from the integrated Seurat object by subsetting the epithelial clusters 3 and 4 (see Sup. Fig. 1A-B). Mesenchymal cells from postnatal timepoints (≥ PND1) were isolated from the integrated Seurat object by subsetting cells based on detectable Vimentin (*Vim*) and *Pdgfra* expression (normalized expression > 0 in the RNA assay). Smooth muscle cells were further defined by first subsetting clusters enriched for *Vim* and *Myh11* expression and subsequently retaining only cells with *Myh11* mRNA within those clusters (normalized expression > 0 in the RNA assay). Gene ontology (GO) enrichment analysis of cluster markers and differentially expressed genes was performed using ShinyGO 0.85^118^. Hallmark gene sets for epithelial and mesenchymal cells were investigated using GSEA^119,120^. Prospective cell trajectories and lineages across pseudotime were predicted with Monocle3^54^ (v1.4.27). Cells were projected into a lower-dimensional space, and trajectory graphs were learned using the reversed graph embedding algorithm. Pseudotime values were computed from a user-defined root node corresponding to the earliest developmental stage (see Fig. 3C) and differentially expressed genes (FDR *P* < 0.01) across the pseudotime trajectory determined.

The 10X Visium HD data consist of FASTQ sequencing files, and a bright-field microscopy image stained with H&E per capture area. The Space Ranger (v.4.0.1) software provided by 10X Genomics was used to align the barcoded spot pattern to the H&E tissue image and to differentiate tissue from background. Data were processed by aligning sequencing reads to the mouse reference genome (refdata-gex-mm10-2020-A). Cell segmentation and downstream spatial analysis and visualization were performed in Seurat, following the recommended workflow for Visium HD data with cell segmentation outputs.

All original datasets will be made publicly accessible (GSE331177 and GSE331354). Source data are provided with this paper. scRNA-seq data used to generate the figures in this paper can also be accessed via our web portal:

For fetal-neonatal uterine atlas: www.genesearch.org/sklab/uterineatlas/

For neonatal endometrial organoids: www.genesearch.org/sklab/neonatalEEO/

For disrupted adenogenesis (E2 and P4): www.genesearch.org/sklab/glanddisruption/

### Mesenchymal-epithelial crosstalk analysis

CellChat^50^ v.2.2.0 was used to infer cell-cell communication using standard pipeline. Briefly, to identify potential interactions, the expression matrix was pre-processed using the in-built functions identifyOverExpressedGenes, identifyOverExpressedInteractions, and projectData with default parameters. Next, computeCommunProb, computeCommunProbPathway, and aggregateNet were used to identify signaling pathways contributing most to outgoing and incoming communication for each cell group. In addition, filterCommunication was used to remove cell-cell communication if there were fewer than 10 cells.

### CellHint label harmonization

Data processing and generation of label harmonization tree were performed as previously described using CellHint^38^. Briefly, the gestational day 4 dataset (GSE275806) from Jia *et al*.^37^ and our fetal-neonatal (E16.5, PND1, PND5, PND12, and PND15) scRNA-seq datasets were concatenated retaining only shared genes (inner join). The combined matrix was library-size normalized (10,000 counts per cell), log-transformed, and processed through batch-aware highly variable gene selection (subset in-place), scaling (max value 10), PCA, and nearest-neighbor graph construction followed by UMAP embedding. Cluster annotations from Jia et al.^37^ were used as input labels for cell type harmonization using *cellhint.harmonize*, specifying the fetal-neonatal dataset as the reference (*dataset_order=[“FetalNeonatal”,“GD4”*]). This returned fine-grained (reannotation) and coarse (group) harmonized cell type labels that were embedded into the *AnnData* object. Supervised data integration was then performed using *cellhint.integrate*, which builds a neighborhood graph constraining neighbor search to transcriptionally similar cell types across datasets. The UMAP was then recomputed on the integrated neighborhood graph. The *cellhint.treeplot* function was used to examine and semi-automatically align the harmonized labels across the two datasets. Concordance between aligned clusters was further validated by comparing the expression of known marker genes.

### Human-mouse cross-species comparison

We leveraged the availability of a scRNA-seq dataset of the developing human fetal uterus (post-conceptional weeks 6 to 20) from Lorenzi *et al.*^14^ to identify conserved transcriptional regulators of epithelial and mesenchymal lineages in mice and humans. To compare transcriptional programs of early uterine development across species, we subsetted our mouse scRNA-seq dataset to mesenchymal and epithelial cell populations and obtained equivalent cell types from first- and second-trimester human Müllerian duct and uterine samples^14^. Gene expression matrices were restricted to one-to-one orthologs between mouse and human, identified using Ensembl BioMart. Highly variable genes (HVGs) were computed independently for each species, and the intersection of the two HVG sets was retained for downstream analysis. To mitigate compositional biases, cells were downsampled within each species so that individual cell types were represented in approximately equal proportions. A low-dimensional embedding was then computed separately for each species by principal component analysis (PCA) on the shared HVG set, preserving species-specific transcriptional structure without forcing alignment into a common embedding space.

Neighborhood graphs were constructed independently on each species’ PCA space using MILO^121^. Cross-species neighborhood matching was performed with scRIMA^39^, which operates agnostically with respect to cell type labels. Briefly, pairwise similarities between mouse and human neighborhoods were calculated using Spearman correlation on mean neighborhood expression profiles across the shared ortholog gene set. The statistical significance of each neighborhood–neighborhood similarity was then assessed, and significant connections were retained to establish cross-species matches. Finally, gene-level conservation was quantified using scRIMA’s Conservation of Patterned Expression (CoPE) score, which identifies genes whose expression patterns are consistently maintained across matched neighborhoods between species.

### Organoid and tissue preparation for immunofluorescence analysis

For tissue processing, female reproductive tracts from 5-15 females were collected per PND/genotype/hormone treatment. Tissues were fixed in 4% paraformaldehyde in PBS at 4C overnight. For EEOs, at least four organoid:Cultrex drops were examined per biological replicate. EEO were incubated in a shaker (30 RPM) with 1 mL of Cell Recovery Solution (Corning, 354253) for 1 h at 4°C. EEO were then fixed using 4% EM grade paraformaldehyde (PFA; Electron Microscopy Sciences, 15710) for 15 min at room temperature, stained with hematoxylin for 10 min, and washed 3-5 times with HBSS supplemented with 10% FBS to remove excess staining. EEOs were resuspended in 60 µL of 2% low-melting agarose (Fisher, BP16525) diluted in HBSS and quickly aliquoted (3-4 drops) on parafilm sheets (3 cm x 3 cm). The agarose droplets containing the EEO were sprayed with 70% ethanol to maintain humidity and incubated for 20 min at RT to let the agarose solidify. EEO drops were carefully transferred to tissue cassettes using #5 forceps. EEO droplets and fixed tissues were dehydrated in ethanol, embedded in paraffin wax, and sectioned (5 µm) as previously described^115^.

### Immunofluorescence staining

Sections were mounted on slides, baked for 30 min at 60 °C, deparaffinized in xylene, and rehydrated in a graded alcohol series. Deparaffinized sections were subjected to antigen retrieval by incubating sections in 1x Tris-EDTA buffer (pH 9.0) at 95 °C for 15 min, followed by cooling to room temperature for 30 min. All slides were blocked with 2% (v/v) normal goat serum (NGS; Invitrogen, 01-6201) in PBS at room temperature for 1 h and incubated with primary antibodies (see Table 3) overnight at 4°C in blocking buffer. Immunofluorescence was performed with Alexa 488, Alexa 594, or Alexa 647-conjugated secondary antibodies (1:500 dilution; Jackson ImmunoResearch, #112-545-143, #111-585-003, #111-605-144). Sections were counterstained with Hoechst 33342 (2 μg/mL; Invitrogen, H3570) before affixing coverslips with ProLong™ Diamond Antifade Mountant (Invitrogen, 36961). Images were taken with a Leica DM6 B upright microscope and Leica K8 camera using Leica Application Suite X (LAS X).

### Quantification and statistical analysis

All mouse and cell culture experiments were performed using at least four biological and technical replicates. The data are presented as the mean ± SEM, as determined from at least three independent experiments. Following the Shapiro-WILK test for normality, the data were analyzed using Student’s t-test or ANOVA with the Bonferroni multiple-comparison test (GraphPad Prism 9). A *p*-value of less than 0.05 was considered statistically significant. Statistical analyses for the genomic experiments were performed using standard statistical tests as described above.

## Acknowledgments

We would like to thank members of the Kelleher and Spencer laboratories for helpful discussions and assistance, and the Genomics Technology Core at the University of Missouri-Columbia. This work was supported by NIH Grant R01HD112315 (A.M.K.) and NIH Grant 1R37HD114609 (T.E.S. and A.M.K.)

## Data and code availability

Sequencing data have been deposited in the Gene Expression Omnibus under accession numbers GSE331177 and GSE331354

